# Effect of aboveground mass removal on toxicity of Geyer’s larkspur, with important implications for grazing management

**DOI:** 10.1101/492900

**Authors:** Kevin E. Jablonski, Paul J. Meiman

## Abstract

The many species of larkspur (*Delphinium* spp. L.) are among the most dangerous poisonous plants on rangelands in the western United States, causing death losses estimated at 2-5% (up to 15%) per year for cattle grazing in larkspur habitat. Other effects, such as altered grazing management practices and consequent lost forage quantity and quality, are significant but poorly understood. Current best management practice recommendations are based on seasonal avoidance, with little evidence that this is practical or effective. Our ongoing research has presented evidence that instead it may be possible to manage grazing such that all cattle eat some larkspur, but none eat a fatal dose. This raises the question of the potential response of larkspur to being grazed. In this study we examine the response of Geyer’s larkspur (*D. geyeri* Greene) to two seasons of 25% or 75% aboveground plant mass removal. The 75% treatment led to significantly lower alkaloid concentrations (mg^•^g^-1^) and pools (mg per plant), while the 25% treatment had a lesser effect. Combined with lessons from previous studies, this indicates that Geyer’s larkspur plants subject to aboveground mass removal such as may occur via grazing can be expected to become significantly less dangerous to cattle. We suggest that the mechanisms for this reduction are both alkaloid removal and reduced belowground root mass, as significant evidence indicates that alkaloids are synthesized and stored in the roots. These results continue to build support for our theory that the solution to the seemingly intractable challenge of larkspur poisoning lies not in avoidance but in the skill of managers and the wisdom of herds.

## Introduction

Larkspur (*Delphinium* spp. L.) are perennial herbaceous plants in the family Ranunculaceae, with approximately 300 total species distributed across the Northern Hemisphere and the mountains of tropical Africa, and 61 species in North America (Warnock, 1995). Larkspur plants constitutively contain numerous norditerpinoid alkaloids, which are potent neuromuscular paralytics capable of damaging or killing many organisms, including humans, mice, insects, and livestock (Welch et al., 2015). In the western United States, researchers have identified eleven species that cause significant cattle death losses, with a recent estimate indicating an average cattle herd loss of 2-5% (15% in some cases) per year, amounting to an estimated $234 million in losses per year in 1988 (Green et al., 2009; Nielsen, 1988; Pfister et al., 1997a; Welch et al., 2015). Such losses have been remarkably intractable for more than a century despite significant research and extension efforts (Cronin and Nielsen, 1972; Glover, 1906; Green et al., 2009; Marsh et al., 1916).

Recommended best management practices in larkspur habitat focus on seasonal avoidance, aimed at reducing exposure to the plants when alkaloid concentration is highest (Pfister et al., 1997b; Welch et al., 2015). Though it is based on sound science, this strategy may create problems of its own as producers lose flexibility to meet their management objectives, both economic and ecological, with little evidence of reduced losses. Regardless of recommendations, many producers abandon grazing on pastures where larkspur is present, either as a precaution or because of past losses (Green et al., 2009). There is a large opportunity cost to this practice, as larkspur tends to grow in resource-rich micro-habitats, where stocker cattle have the potential to gain 2.5 pounds per day (Green et al., 2009; Pfister et al., 1997a). Other producers appear to accept the risk of deaths, experiencing gains when lucky and losses when not. All told, larkspur presents one of the most significant challenges to grazing management in the western US.

One potential alternative to current avoidance-based strategies is to manage grazing such that all individuals consume some larkspur, but no individual consumes a lethal dose. Our recent paper (Jablonski et al., 2018) presented an agent-based model simulation that indicated this may be possible if cattle are managed for high stocking density, high herd cohesion, or both. The potential for this solution has also been noted anecdotally by producers (e.g. Smith et al., 2010). This raises questions about the potential response of larkspur to being grazed, as well as the effects of years of generally not being grazed by cattle.

Two previous studies have examined the response of larkspur species to manual clipping. Laycock (1975) tested the effect of clipping on groups (n= 11-16) of duncecap larkspur (*D. occidentale* S. Watson) in Idaho and found a reduction in both alkaloid concentration and plant mass in subsequent years. Ralphs and Gardner (2001) tested the effect of clipping on twenty subalpine larkspur (*D. barbeyi* Huth) plants and found a reduction in plant mass but not in alkaloid concentration in subsequent years. Both studies consisted of a single-level treatment of clipping of the full plant at or near ground level, and both interpreted their findings through the lens of mechanical clipping. Neither mentioned the potential long-term ramifications of non-grazing of larkspur on plant vigor and toxicity nor the potential for grazing to play a role similar to mechanical clipping.

Nearly all previous studies of larkspur toxicity have focused on alkaloid concentration (typically mg^•^g^-1^) as the key measure of poisoning risk to cattle. However, we have come to focus on aboveground alkaloid pool as a more useful comprehensive measure of poisoning risk. Aboveground alkaloid pool is the product of aboveground plant biomass and alkaloid concentration, and can be measured on a per-plant, per-hectare, or per-pasture basis. An understanding of aboveground alkaloid pools is essential if we shift from thinking of larkspur consumption as something to be avoided to something that can be managed at sub-lethal levels.

We focused on larkspur measurements at the bud stage of growth. We chose bud stage because this is when aboveground alkaloid pools are maximized (Ralphs and Gardner, 2003). This growth stage of larkspur also corresponds to the one of the most favorable times for livestock grazing in the foothill rangelands of *D. geyeri* Greene. It is also a time of year when otherwise attractive pastures are often avoided when larkspur is abundant (Green et al., 2009).

In this paper we assess the potential response of larkspur to incomplete removal of aboveground plant material, as might occur via grazing. Specifically, we compare the effect of two seasons of 25% and 75% removal of aboveground plant mass to unclipped (control) plants. In combination with our agent-based model findings, the results continue to build support for an alternative approach to one of the most intractable challenges faced by western livestock producers.

## Methods

### Study area

We collected field data for *D. geyeri* in June of 2016, 2017, and 2018 at the Colorado State University Research Foundation Maxwell Ranch, a working cattle ranch in the Laramie Foothills ecoregion of north-central Colorado that is a transition zone between the Rocky Mountains and the Great Plains. There are significant populations of Geyer’s larkspur at several locations on the ranch which have historically created management challenges and resulted in poisoned cattle. We focused our sampling within established research plots in an area of particularly dense Geyer’s larkspur stands (N40° 54.85’ W105° 13.64’).

### Aboveground dry mass estimation

Because repeated measurements of individual plants were required, we devised a non-destructive method of estimating aboveground Geyer’s larkspur plant dry mass (Catchpole and Wheeler, 1992). While challenging for herbaceous plants, such a method can be reliable if it is applied to an individual species, is calibrated for the specific situation, and incorporates multiple plant traits (Catchpole and Wheeler, 1992; Ohsowski et al., 2016). In this case a linear model was sufficient.

To create the model, in 2016 we randomly selected 120 Geyer’s larkspur plants of any size once all plants had reached early bud stage. We began with number of leaves, number of stems, and total stem length as predictor variables for plant mass, chosen for their ease of measurement and hypothesized correlation with mass. To measure number of leaves, we simply counted the total number of fully-formed leaves on each plant. We did not include leaves that were mostly brown and dead but did include leaves that were mostly green with some browning. To measure number of stems, we counted the total number of stems emerging from the ground for an individual plant. To measure total stem length (cm), we summed the total length of all the stems for a given plant. We then cut, dried, and weighed each plant

We analyzed the mass estimation data using R statistical software, version 3.5.1 (R Core Team, 2018). We used multiple linear regression within an information-theoretic framework (R package MuMIn) to generate and compare models (Anderson, 2008). Because the raw data violated assumptions of homoscedasticity and normal distribution of error, we performed a natural log transformation of the predictor variables and plant dry mass. A variance inflation factor test then indicated that there was a problem with multicollinearity among the predictor variables (Graham, 2003). This was largely driven by a strong correlation between number of stems and total stem length. Not wanting to drop one of the terms and lose the unique information it provided, we instead combined these two terms into one term, length per stem. During our field work, we had in fact suspected that length per stem would be a good predictor of plant mass, as it corrected for cases where there were many short stems. This change solved the issue of multicollinearity and greatly lowered the standard error on the coefficient estimates. We used corrected AIC to assess the whether the full model was superior to models containing each of the single predictor variables.

### Mass removal treatments

To apply the mass removal treatments, we began in June 2016 by randomly selecting 81 *D. geyeri* plants and marking them with a durable metal stake. We then assigned them to one of three treatments: unclipped control, 25%, or 75%. Individual plants remained in their assigned treatment category for the duration of the study.

In June of both 2016 and 2017, once all plants were in the bud stage, we recorded number of leaves and length per stem. We then applied the appropriate treatment to the plant. For the 25% and 75% mass removal treatments, we used our measured variables to estimate the appropriate amount of plant material to remove. For the control group, we removed three leaves for assessment of alkaloid concentration but otherwise left the plant untreated. In June of 2018 we measured the same variables and then cut each entire plant, regardless of treatment level, to compare our linear model mass predictions to the measured actual mass.

### Sample preparation and alkaloid analysis

All plant samples were dried, weighed, and ground until sufficiently fine to pass through a 2 mm mesh screen. They were then individually labeled and shipped to the USDA Poisonous Plant Laboratory in Logan, UT for assessment of alkaloid concentration. The dried ground samples were then extracted and analyzed using standardized methods for alkaloid assessment in larkspur (Gardner et al., 1999). For this study, we focus on the reported measures for total MSAL alkaloid concentration in the plants.

### Treatment data analysis

Note that we used actual weights instead of predicted weights for 2018. Group means were similar, but we saw no reason to use predicted weights when we had access to actual weights. Due to concerns about non-normal distribution of error, heteroscedasticity, and difficulty of transforming percent change data, we used a Steel-Dwass non-parametric multiple comparisons test to examine differences in the distribution of data among the three treatments (Douglas and Michael, 1991). The Steel-Dwass test is fairly conservative and thus helps avoid Type-I (false-positive) errors. We used JMP 13.0.0 (SAS Institute, 2016) for analysis and data visualization.

## Results

### Aboveground mass estimation

Data for the linear model using number of leaves (*leaves*) and length per stem (*lpstem*) are shown in Table 1. AIC_c_ scores indicated that the model containing both *leaves* and *lpstem* was best, with single-factor models highly implausible in comparison, given the data (Anderson, 2008). It is worth noting that in 2018, when we obtained estimated and actual dry mass for all plants, the estimated mean for all plants was 2.34 g while the actual weighed mean was 2.36 g. This demonstrates that even if standard error is relatively high a moderate number of samples will ensure accuracy to the group mean.

**Table 1.** Results for multiple linear regression of natural log transformed data for predicting *D. geyeri* plant mass from plant characteristics (n=120).

| Variable (ln transf.) | B<sup>1</sup> | SE | CI<sup>2</sup> |
| --- | --- | --- | --- |
| Intercept | -4.90 | 0.19 | -5.28, -4.53 |
| *leaves* | 1.09 | 0.04 | 1.01, 1.18 |
| *lpstem* | 0.79 | 0.07 | 0.65, 0.92 |
| SE of residuals= 0.29 |  |  |  |
| Adj. R<sup>2</sup>= 0.92 |  |  |  |

<sup>1</sup>All coefficients significant p<0.001 <sup>2</sup>95% confidence interval

### Mass removal treatments

Of the 81 larkspur plants initially selected, we could not relocate the markers for four during either the second or third season. We attribute losing these markers to rodent activity. Of the remaining 77 plants, three died after 2017 but prior to 2018. Of these, one was in the 25% mass removal treatment group and two were in the 75% mass removal treatment group. As each of these three plants had declined greatly when measured in the second year, we attributed these deaths to the treatments and thus included them in the analysis. Ultimately, there were 26 plants in the control group, 25 plants in the 25% group, and 26 plants in the 75% group.

It is first important to note that the all three treatment groups experienced overall declines in mass, MSAL alkaloid concentration, and MSAL alkaloid pools (Fig 1). We attribute this general decline (which we also observed, anecdotally, across the study ranch) to a return to relatively normal precipitation patterns from heavier than normal precipitation in 2013-2015. Despite this general decline, there were nevertheless clear differences among the treatments.

**Fig 1.**
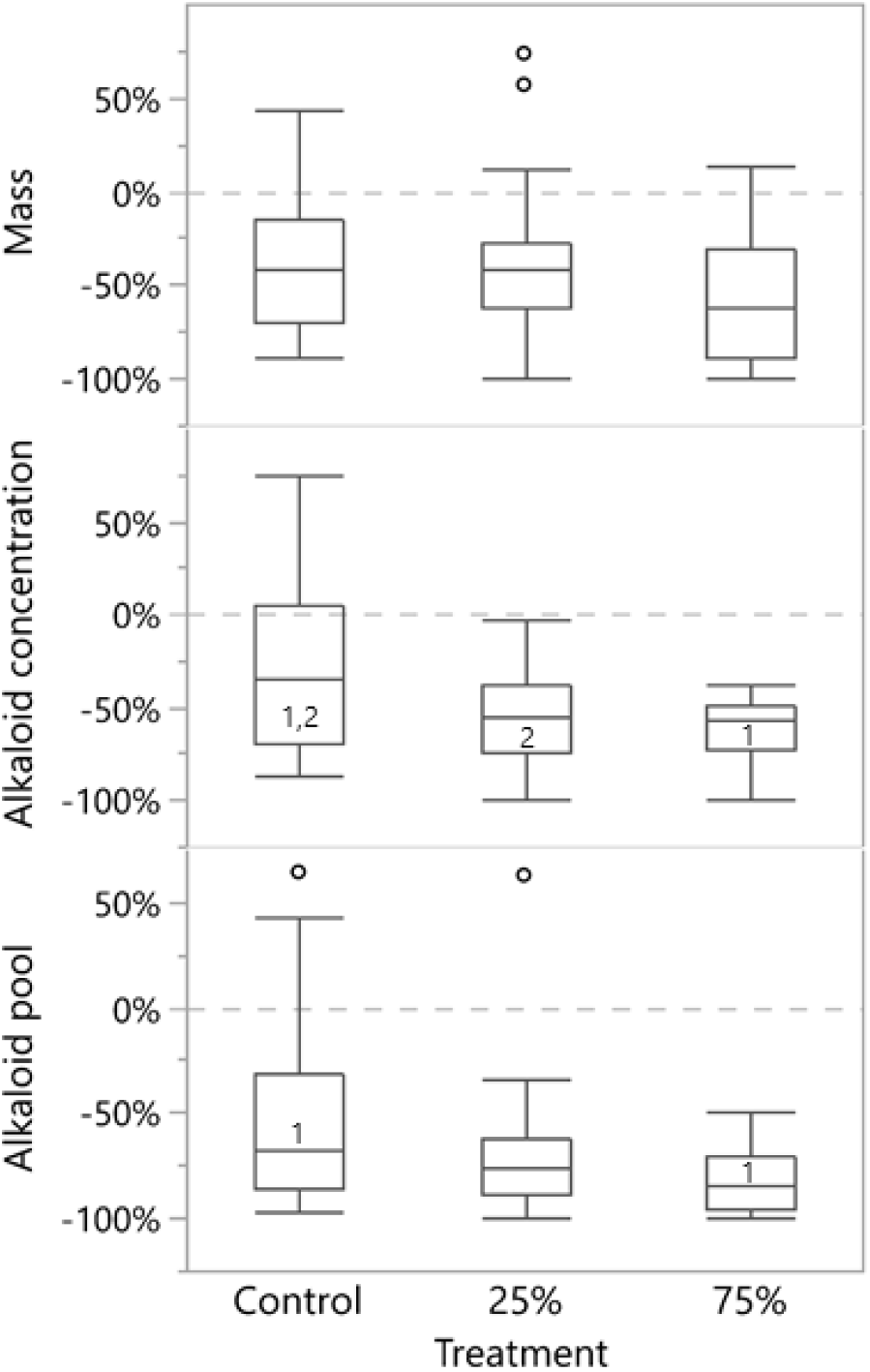
Box plots (with outliers) of the distribution of *D. geyeri* treatment responses (2016-2018), as measured by percent of 2016 plant mass, MSAL alkaloid concentration, and MSAL alkaloid pools. Shared numbers indicate a significant difference in group medians (1-indicates that the group medians were significantly different at α=0.05. 2-indicates that the group medians were significantly different at α=0.1).

While there was some indication of a decline in mass due to treatments, these differences were not statistically significant. On the other hand, significant declines in MSAL alkaloid concentration and plant alkaloid pools occurred at the 75% treatment level versus the control and, for concentration, at both treatment levels. Additionally, the treatments had a clear effect on the variance of alkaloid concentration and alkaloid pools. Table 2 shows the mean values for the responses, as well as the measured morphological changes.

**Table 2.** Mean *D.geyeri* treatment responses (2016-2018) for plant mass, MSAL alkaloid concentration, MSAL alkaloid pool, number of leaves, total stem length, and number of stems.

|  | Percent change |  |  |  |  |  |
| --- | --- | --- | --- | --- | --- | --- |
|  | Mass | Conc. | Pool | Leaves | Stem length | Stems |
| Control | -36.9 | -25.6 | -53.9 | -16.9 | -34.4 | 8.5 |
| 25% | -38.9 | -54.3 | -70.4 | -29.3 | -36.2 | -9.7 |
| 75% | -55.3 | -61.1 | -81.7 | -42.0 | -47.6 | -28.9 |

An examination of year-on-year results reveals that plant mass decreased greatly after the first year, while alkaloid concentration responded more strongly after the second year (Fig 2). Alkaloid pools, as the product of mass and concentration, responded more evenly. Note the increase in control group alkaloid concentration in 2017.

**Fig 2.**
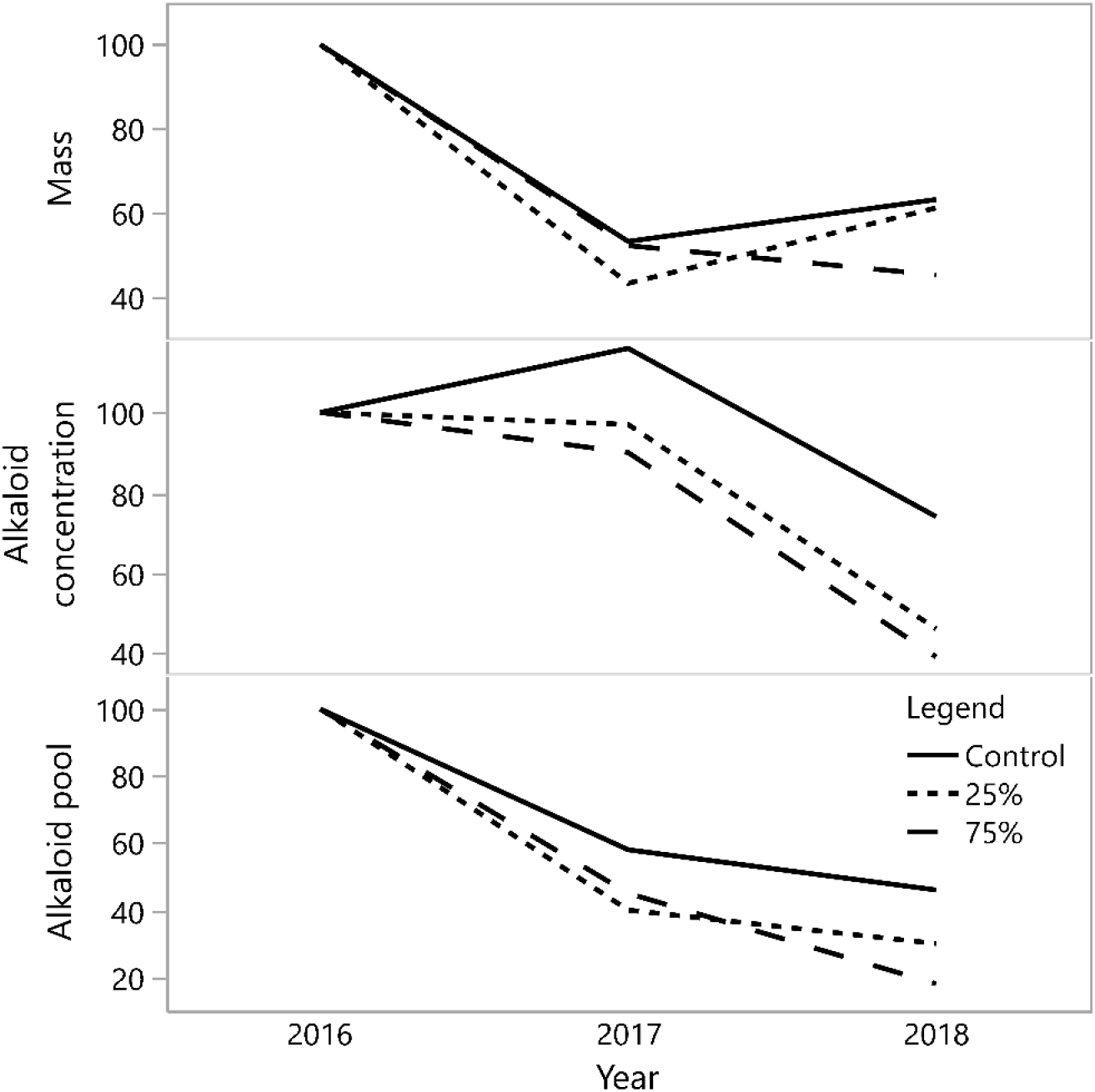
Indexed (2016=100) mean *D. geyeri* treatment responses for plant mass, MSAL alkaloid concentration, and MSAL alkaloid pools by year.

## Discussion

The mechanisms by which larkspur plants generate, transfer, and store toxic alkaloids are still somewhat unclear, and may differ among species. However, the most likely general scenario is as follows: Alkaloids are synthesized in the roots and translocated upward into stems and leaves early in the growing season (Ralphs and Gardner, 2003). This relatively fixed amount of alkaloids is then diluted as aboveground plant mass increases during the growing season (Gardner and Pfister, 2007; Ralphs et al., 2000). As the plant begins to senesce, most of the alkaloids are translocated back to the roots and stored for the following growing season (Ralphs et al., 2000; Ralphs and Gardner, 2003).

Norditerpenoid alkaloids are high in nitrogen and are thus expensive for larkspur plants to produce. Nevertheless, numerous studies have indicated that these alkaloids are generated constitutively and that levels are relatively unresponsive to stress or environmental factors, being frequently measured at a concentration that is higher than necessary for general plant defense. These are unusual dynamics and beg further exploration.

Because insect pressure on larkspur is quite low and the different species are generally quite nutritious for large herbivores (Pfister et al., 1997c), the place to start this exploration is with herbivore pressure. While science has been studying larkspur in the western US for just over 100 years, the genus *Delphinium* has co-evolved with herbivores for millions of years (Jabbour and Renner, 2012). We suggest that this most recent 100-year period is highly anomalous in that humans have, broadly-speaking, replaced a diverse set of millions of wild large herbivores with a similar number of domestic large herbivores (Burkhardt, 1996) and then generally managed cattle to avoid grazing larkspur during its growing season (Green et al., 2009). Released from grazing pressure, larkspur has continued to produce alkaloids at its historical rate, but these alkaloids are no longer being consistently removed from the plants via herbivory. Meanwhile, larkspur has also increased in abundance and mass, as is suggested by its plant characteristics (i.e., tall perennial forb growing in a dry climate), which place it into the category of a likely “decreaser” under heavy grazing (Diaz et al., 2007; Milchunas et al., 1988).

The combination of Laycock (1975), Ralphs and Gardner (2001), and this study indicate that larkspur plants that suffer moderate-to-heavy grazing will indeed contain lower amounts of toxic alkaloids, greatly reducing the risk to grazing herbivores. Conversely, ungrazed larkspur will hold onto precious alkaloids, increase its root mass, increase its alkaloid synthesis and storage capacity (Ralphs et al., 2000; Ralphs and Gardner, 2003), and increase the amount and toxicity of the plant material that it presents to herbivores. In short, our management of cattle to avoid grazing larkspur, especially when it is most vulnerable, may have worsened, if not created, the problem of consistent cattle poisoning.

There is of course complexity and nuance to this story. For example, while both previous studies found a significant difference in mass after full aboveground plant mass removal, we found a lesser (p=0.16) effect from 75% aboveground mass removal. Whether this was due to the lower level of mass removal, environmental conditions lowering overall plant mass regardless of treatment, or something else, was unclear. Because we saw a continued decline in mass in 2018 from the 75% removal treatment while the other two treatments increased in mass (Fig 2), we suspect that environmental conditions were the driver in 2017.

Interestingly, we saw a lag between a reduction in aboveground plant mass and a reduction in alkaloid concentration (Fig 2). If we consider root mass and vigor to be the driver of alkaloid concentration, this makes perfect sense. A significant removal of aboveground biomass in one year would likely lead to reduced root mass the following year, and thus reduced capacity to synthesize and store toxic alkaloids. However, we believe that alkaloid removal also explains the especially strong decline in concentration across the two mass removal treatment levels. By simply removing more alkaloids from the plant than it may be able to regenerate in a year, mass removal lowers concentration. This means that grazing may have a double effect on larkspur toxicity—it steals what was expensive to produce and also makes it harder to re-produce it.

If we accept the control group as status quo under normal fluctuations, even when the status quo alkaloid pool declined by greater than 50%, removal of 25% of aboveground larkspur mass led to an additional mean yearly decline of 8.2% in alkaloid pools while removal of 75% of aboveground mass led to an additional mean yearly decline of 13.9% in alkaloid pools. Though it is impossible to accurately extrapolate further, with the cumulative declines among living plants and expected continued deaths of weaker plants (3/51 treated plants died from the treatments), we think it is reasonable to expect a 50% decline in pasture-level alkaloid pools within a few years of moderate-to-heavy grazing during bud stage. Because it is believed that the majority of larkspur deaths occur from brief periods of over-ingestion (Pfister et al., 1997c), cutting alkaloid pools in half would be likely to dilute risk below a threshold where the animal satiates on larkspur before they consume a lethal dose of alkaloids. Indeed, the data for our control group demonstrate that alkaloid pools fluctuate widely even without intervention, providing explanation for such a threshold as a driver of inconsistent losses among producers.

## Conclusions and implications

Grazing management in the broad array of environments that constitute larkspur habitat is always a complex, multifaceted endeavor. This means that any solution to the seemingly intractable challenge of larkspur poisoning must account for the spatiotemporally unique multiple objectives that producers and managers seek to fulfill on grazing lands, rather than treating larkspur as an isolated, singular challenge. Thus we offer no such solution here. Instead, this study continues to build support for our theory that the solution to the larkspur challenge lies not in avoidance but rather in the skill of managers and the wisdom of herds.

Building on the many years of research on larkspur toxicology patterns, we have demonstrated that Geyer’s larkspur plants subject to aboveground mass removal similar to what might occur with grazing can be expected to become significantly less dangerous to cattle. When we consider what is known about alkaloid synthesis, translocation, and storage in larkspur, the reasons for this are clear. We can expect that *D. geyeri* and other larkspur species will be “decreasers” and, as such, grazing will lead to reduced above- and below-ground biomass, with consequent reductions in alkaloid concentration and pools.

For managers, we echo the advice from Jablonski et al. (2018) of caution in the face of this threat, especially for those producers that have experienced serious losses. Nevertheless, these two papers indicate that amid high populations of *D. geyeri* it is possible to manage grazing such that all cattle consume larkspur but none die and that, having done so, the risk will decline over time. This may require alterations to management practices and increased attention to herding, but we expect that improved management flexibility and increased capacity to graze pastures when they are at full productivity will offset these increased efforts. Avoidance may always be simpler, but we put faith in the fact that ranchers prefer effectiveness to ease.

## Acknowledgements

We are grateful to Dale Gardner of the USDA Poisonous Plants Laboratory in Logan, UT for his generous assistance in testing alkaloid concentrations. We also thank Joel Vaad, Manager of the Colorado State University Research Foundation Maxwell Ranch, for providing space for the research and for sharing his thoughts and time. Tanner Marshall assisted in study design, data collection, and sample preparation. This work was supported by funds from the Colorado State University Agricultural Experiment Station [grant number AES-0650] and the Colorado State University Research Foundation.

